# Representativeness and robustness of TCR repertoire diversity assessment by high-throughput sequencing

**DOI:** 10.1101/242024

**Authors:** Wahiba Chaara, Encarnita Mariotti-Ferrandiz, Ariadna Gonzalez-Tort, Laura Florez, Adrien Six, David Klatzmann

**Author notes:** equal contribution (Wahiba Chaara, Encarnita Mariotti-Ferrandiz). **Authors’ contributions:** WC and EMF equally contributed to the work (^$^).WC performed all the bioinformatics analyses. AG and LF prepared the samples. WC, EMF, AS and DK conceived the studies, designed the experiments and analysed the results. WC, EMF, AS and DK wrote the first draft of the manuscript, with input from all authors. DK initiated and obtained funding for the study. LF was funded by a “DIM Région Ile-de-France” doctoral fellowship. The work of WC, EMF, AS and DK is funded by the Assistance Publique-Hôpitaux de Paris, INSERM, and Sorbonne Universités UPMC (Paris 6). The study is part of the LabEx Transimmunom (ANR-11-IDEX-0004-02) and ERC Advanced Grant TRiPoD (322856) funding obtained by DK. The RNA sequences presented in this study have been submitted to Sequence Read Archive (SRA; https://www.ncbi.nlm.nih.gov/sra) as Bioproject PRJNA408306, under accession numbers SRR6068973, SRR6068972 and SRR6068975 (Biosample SAMN07682929) and SRR6068974, SRR6068969, SRR6068968, SRR6068971, SRR6068970, SRR6068967, SRR6068966 and SRR6068976 (Biosample SAMN07682930).

## Abstract

High-throughput sequencing (HTS) has the potential to decipher the diversity of T cell repertoires and their dynamics during immune responses. Applied to T cell subsets such as T effector and T regulatory cells, it should help identify novel biomarkers of diseases. However, given the extreme diversity of TCR repertoires, understanding how the sequencing conditions, including cell numbers, biological and technical sampling and sequencing depth, impact the experimental outcome is critical to properly use of these data. Here we assessed the representativeness and robustness of TCR repertoire diversity assessment according to experimental conditions. By comparative analyses of experimental datasets and computer simulations, we found that (i) for small samples, the number of clonotypes recovered is often higher than the number of cells per sample, even after removing the singletons; (ii) high sequencing depth for small samples alters the clonotype distributions, which can be corrected by filtering the datasets using Shannon entropy as a threshold; (iii) a single sequencing run at high depth does not ensure a good coverage of the clonotype richness in highly polyclonal populations, which can be better covered using multiple sequencing. Altogether, our results warrant better understanding and awareness of the limitation of TCR diversity analyses by HTS and justify the development of novel computational tools for improved modelling of the highly complex nature of TCR repertoires.

## INTRODUCTION

Understanding the specificity of T cells involved in immune responses is of utmost importance in many fields of immunology. T cells are characterised by the expression a unique T cell receptor (TR), which is clonally generated by somatic rearrangement of the V, D and J genes belonging to the TR genomic locus during thymic T cell differentiation (Davis and Bjorkman, 1988). This process leads to the generation of a huge diversity of TR, defining a repertoire of antigen recognition, the hallmark of the adaptive immune response. Immunoscope analysis (also called CDR3 spectratyping) has long been the standard technique for TR repertoire analyses (Boudinot et al., 2008). Although Immunoscope analysis has been very useful, it misses the key parameters of TR diversity, which include nucleotide sequence, codon usage, and amino acid composition. High-throughput sequencing (HTS) of the adaptive immune receptor rearrangements (RepSeq) expressed in a lymphocyte population now overcomes previous limitations, providing a thorough and multifaceted measure of diversity (Six et al., 2013). Several studies have already highlighted the feasibility of HTS for the analysis of TR repertoire diversity in various immune contexts (Bergot et al., 2015; Dash et al., 2017; Dong et al., 2017; Freeman et al., 2009; Glanville et al., 2017; Kuang et al., 2017; Langerak et al., 2017; Maceiras et al., 2017; Madi et al., 2017; Marrero et al., 2013; Poschke et al., 2016; Sims et al., 2016; Thapa et al., 2015; Thomas et al., 2014). However, while the amount of information and the depth of analysis provided by this technique are unprecedented, the representativeness and robustness of the data obtained remain to be established.

RepSeq is a numbers game (Benichou et al., 2012), particularly dependent on sequencing depth and therefore on sampling. When monitoring T cell leukaemia or highly expanded antigen-specific TCRs following an infection, the sampling and depth of sequencing might not be critical parameters. But things are different when studying TR repertoire diversity in physiological conditions, when describing the basics of immune repertoire generation and selection or in immune contexts where subtle or qualitative modifications may be involved in the pathophysiological outcome, such as in complex infectious diseases (Emerson et al., 2017; Heather et al., 2016; Mariotti-Ferrandiz et al., 2016), autoimmune disorders (Madi et al., 2014; Marrero et al., 2013, 2016; Pugliese, 2017; Rossetti et al., 2017; Seay et al., 2016; Zhao et al., 2016) and transplantation follow-up (van Heijst et al., 2013; Lai et al., 2016; Theil et al., 2017). However, RepSeq necessarily implies sampling: i) only a fraction of the cells from peripheral blood or an organ (or a fragment of that organ in humans) is harvested; ii) only a fraction of the RNA/DNA extracted from these cells is used for sample preparation; and finally, iii) only a fraction of the library is used for a sequencing run. These different levels of experimental sampling are likely to affect the observed diversity.

This is a genuine issue described in ecology studies, as *“the absence of observation of a species can be either real or the effect of a subsampling”* (Magurran, 2004). Previous studies showed that the number of clonotypes observed is positively correlated with sampling size (Madi et al., 2014; Robins et al., 2010; Shugay et al., 2013). This is important, as studies performed in humans are mostly based on peripheral blood, a compartment that represents only around 2% of the total T lymphocyte population. Warren et al. (Warren et al., 2009) compared TR repertoires from two blood samples from the same individual and found a limited number of shared clonotypes (^~^ 10%). They concluded that a considerable proportion of the peripheral blood TR repertoire is unseen when observed randomly (Fisher et al., 1943; Warren et al., 2009).

The depth of the sequencing is another confounding factor for TR repertoire diversity studies, since an insufficient number of sequences produced would not adequately assess the molecular diversity of the sample analysed. To ensure the statistical representativeness of the data produced with regards to the population of interest, two rules should be considered (Greiff et al., 2015a): i) the number of sequences produced must be at least equivalent to the clonal richness of the population of interest; ii) the rarer a clone, the greater the sequencing depth needed to detect it. Therefore, the RepSeq strategy must be adapted to the nature of the samples and the biological questions investigated (Bashford-Rogers et al., 2014).

While most studies seek to assess the similarity between the TR repertoires of several samples, without any knowledge of what level of similarity can be observed at best, it seems crucial to determine the limits of this approach in order to be able to interpret the data properly. In this study, we first investigated the impact of the depth of sequencing, in relation to the size of the population analysed, on the observed TR repertoire diversity. We found that a small sample size is negatively affected by a too high, yet average in common practice, sequencing depth, and proposed an analytical approach to recover the “true” repertoire diversity. We then questioned the representativeness of a single RepSeq experiment by multiple sequencing of the same sample and demonstrated that performing a single sequencing run, even at high depth of sequencing, does not allow exhaustive observation of the existing clones in a polyclonal population. Finally, we addressed these experimental biases by computational simulation on RepSeq data reflecting several levels of clonality and sequencing depth, to have a better assessment of the robustness of the experimental observations.

## MATERIALS & METHODS

### Mice

Eight- to 12-week-old female Balb/C Foxp3-gfp (C.129X1-Foxp3tm3Tch/J) and 24-to 26-week-old male C57Bl/6 Foxp3-GFP mice, both expressing the green fluorescent protein (GFP) under the promoter of Foxp3 gene, were, respectively, provided by V. Kuchroo, Brigham and Women’s Hospital, Boston, MA and B. Malissen of the Centre d’Immunologie de Marseille Luminy (France). All animals were maintained in University Pierre and Marie Curie Centre d’Expérimentation Fonctionnelle animal facility under specific pathogen-free conditions in agreement with current European legislation on animal care, housing and scientific experimentation (agreement number A751315). All procedures were approved by the local animal ethics committee.

### Cell preparation

Fresh total cells from spleen were isolated in PBS1X 3% foetal calf serum (FCS) and stained for 20 min at 4°C with anti-Ter-119-biotin, anti-CD11c-biotin and anti-B220-biotin antibodies followed by anti-biotin magnetic beads (Miltenyi Biotec) labelling for 15 min at 4°C. B cells and erythrocytes were depleted on an AutoMACS separator (Miltenyi Biotec) following the manufacturer’s procedure. Enriched T cells were stained with anti-CD3 APC, anti-CD4 Horizon V500, anti-CD8 Alexa 700, anti-CD44 PE and anti-CD62L efluor 450. 6.10^5^ CD3+CD4+GFP-Teff cells were sorted on a BD FACSAria II (BD Biosciences, San Jose, CA) with a purity > 99%. Sorted cells were stored in Trizol (Invitrogen) or RNAAquous (Ambion, Inc/Life Technologies, Grand Island, NY, USA) lysis buffer.

### TR library preparation

RNA was extracted following the manufacturer’s recommendations and cDNA synthesis was performed with the Qiagen OneStep RT-PCR kit (Qiagen Inc., Valencia, CA, USA) and mouse T cell beta receptor (MTBR) primers provided with the mouse iR-Profile kit (iRepertoire Inc., Huntsville, AL, USA). cDNA was amplified by two rounds of PCR according to the manufacturer’s recommendations. The TR beta library was sequenced using Illumina on a MiSeqv2 kit.

### RepSeq data processing

#### Data annotation

The RepSeq fastq files were demultiplexed by iRepertoire Inc. and then annotated using clonotypeR (Plessy et al., 2015) to identify productive TRB sequences. Clonotypes were defined as unique combinations of TRBV-CDR3-TRBJ segments.

#### Sequencing error correction

Annotated sequences were clustered per TRBV-TRBJ combination and similar clonotypes collapsed as follows: Within each TRBV-TRBJ cluster, the clonotypes observed once (singletons) were separated from the others to constitute two groups. A Levenshtein distance was then calculated between the CDR3 peptide sequences of each clonotype of the two groups. The Levenshtein distance (lev) is a string metric measuring the minimum number of single-character edits (insertions, deletions or substitutions) required to change one sequence into another (Levenshtein, 1966).

When comparing the CDR3 peptide sequences of singleton with that of a “non-singleton” sequences, if lev_seq1_,_seq2_=1, their respective nucleotide sequences are then compared. If the two corresponding nucleotide sequences are also distant by 1, the singleton is considered as erroneous and considered as the “non-singleton” clonotype.

#### Dataset normalisation

Using the function *rrarefy* from the Vegan R package (Oksanen et al., 2013), randomly rarefied datasets were generated to given sample size. The random rarefaction was made without replacement.

### Diversity profiles

Rényi entropy is a generalisation of Shannon entropy, initially developed for information theory. We applied this mathematical function to clonotype frequencies to assess their diversity within each dataset. Rényi entropy is function of a parameter α, a strictly positive real number that differs from 1 and allows the definition of a family of diversity metrics spanning from (i) the species richness (α=0), which corresponds to the number of clonotypes regardless of their abundance to (ii) the clonal dominance (α → +∞), corresponding to the frequency of the most predominant clonotype. For α =1, the Shannon diversity index is computed. The exponential of the Rényi entropy defines a generalised class of diversity indices called Hill diversities, which can be interpreted as the effective number of clonotypes in the datasets (Hill, 1973) and thereby is used to build a diversity profile.

### RepSeq simulation algorithm

A. 2.10^6^ clonotype library construction with the tcR package

Based on the estimated total number of clonotypes in a mouse, a 2.10^6^ TRB CDR3 library was generated with the tcR package following the probability rules of V(D)J rearrangement established in Murugan et al. (Murugan et al., 2012):

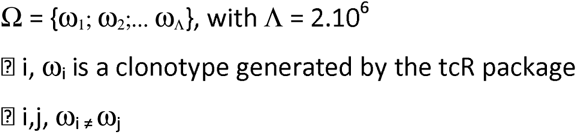

B. Construction of 6.10^5^ sequence datasets following particular Zipf distributions

Based on the demonstration by Greiff et al. (41) that clonotype frequencies determined from RepSeq datasets generally follow a Zipf distribution with a particular *α* ∊ [0, 1] parameter, we chose to use the Zipf-Mandelbrot law implemented in the zipfR R package (Evert and Baroni, 2007) to simulate clonotype distributions. The probability density function used for simulations is given by

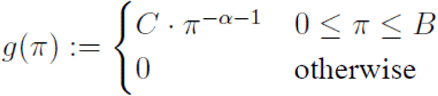

with two free parameters: *α* ∊ [0, 1] and B ∊ [0, 1] and a normalising constant C. B corresponds to the probability π1 of the most frequent species (clonotype).

Seven Zipf distributions were generated with the following Zipf parameters:

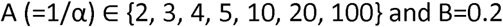

For each Zipf parameter combination, a list Z_A_ is randomly generated as follows:

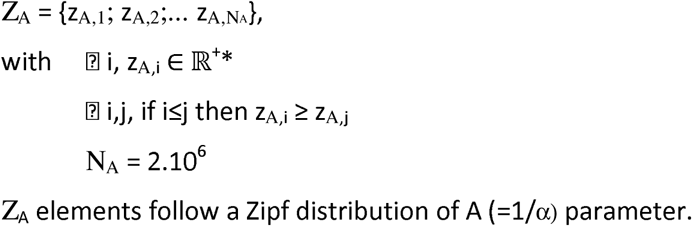

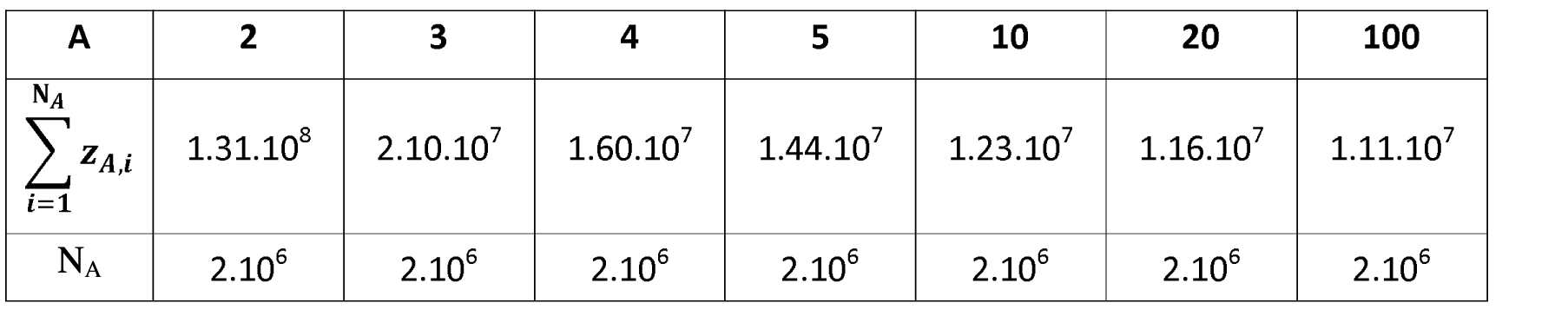

C. For each A parameter, the 2.10^6^ Z_A_ values were randomly assigned to the clonotype collection to obtain seven TRB clonotype repertoires.

D. To obtain the final seven datasets, each of them were rarefied using the function *rrarefy* from Vegan R package to a 6.10^5^ size datasets.

#### Rarefaction at increasing sizes

Each of the seven simulated datasets were rarefied into a series of six datasets of size D ∊ {500, 100, 5000, 5.10^4^, 5.10^s^, 1.10^6^}. For each value of D, subsamples of TRB sequences were randomly produced using the *vegan::rrarefy* function (without replacement). This process was iteratively repeated 100 times with replacement. For each resulting series of subsamples, clonotype counts were calculated and used to assess the median and 95% CI values of Morisita-Horn index (MH; (Horn, 1966)) between them and the original dataset (representativeness) and between each other (robustness).

Subsample compositions were also compared to evaluate the level of overlap between 3 subsamples according to the dataset size.

For each D, combinations of 3 Z_A_ dataset subsamples were randomly selected to determine the proportion of clonotypes observed once, twice or in the 3 subsamples. This process was performed 100 times to calculate the median and 95% CI of each result.

Since the 95% CI values obtained for MH and overlap proportion were similar to the medians, they are not indicated in the corresponding figures and tables.

## RESULTS

### Impact of sequencing depth on the representativeness of the repertoire diversity

With advances in HTS technologies, a million sequences per sample has often become the minimum number of outputs in RepSeq studies. Besides, small samples are often studied. Thus, to determine the minimum number of sequences required for a representative repertoire, we first explored how the number of raw reads could affects the repertoire description according to the sample size. We chose to analyse a mouse sample with high diversity and used the CD4+Foxp3-cell population (Teffs) previously described as very diverse (Bergot et al., 2015). 6.10^5^ CD4+GFP-Teff cells from female Balb/C Foxp3<GFP= splenocytes where sorted. RNA was extracted from these cells and diluted in order to obtain aliquots containing the RNA amount equivalent to what would be obtained from 50 000, 5 000, 1 000 or 500 cells (Figure 1A). Two replicates per dilution were prepared. For simplicity in the text, the sample size will be defined according to the theoretical equivalent cell number for each aliquot. Sequencing was performed on RNA amplified by multiplex PCR.

**Figure 1:**
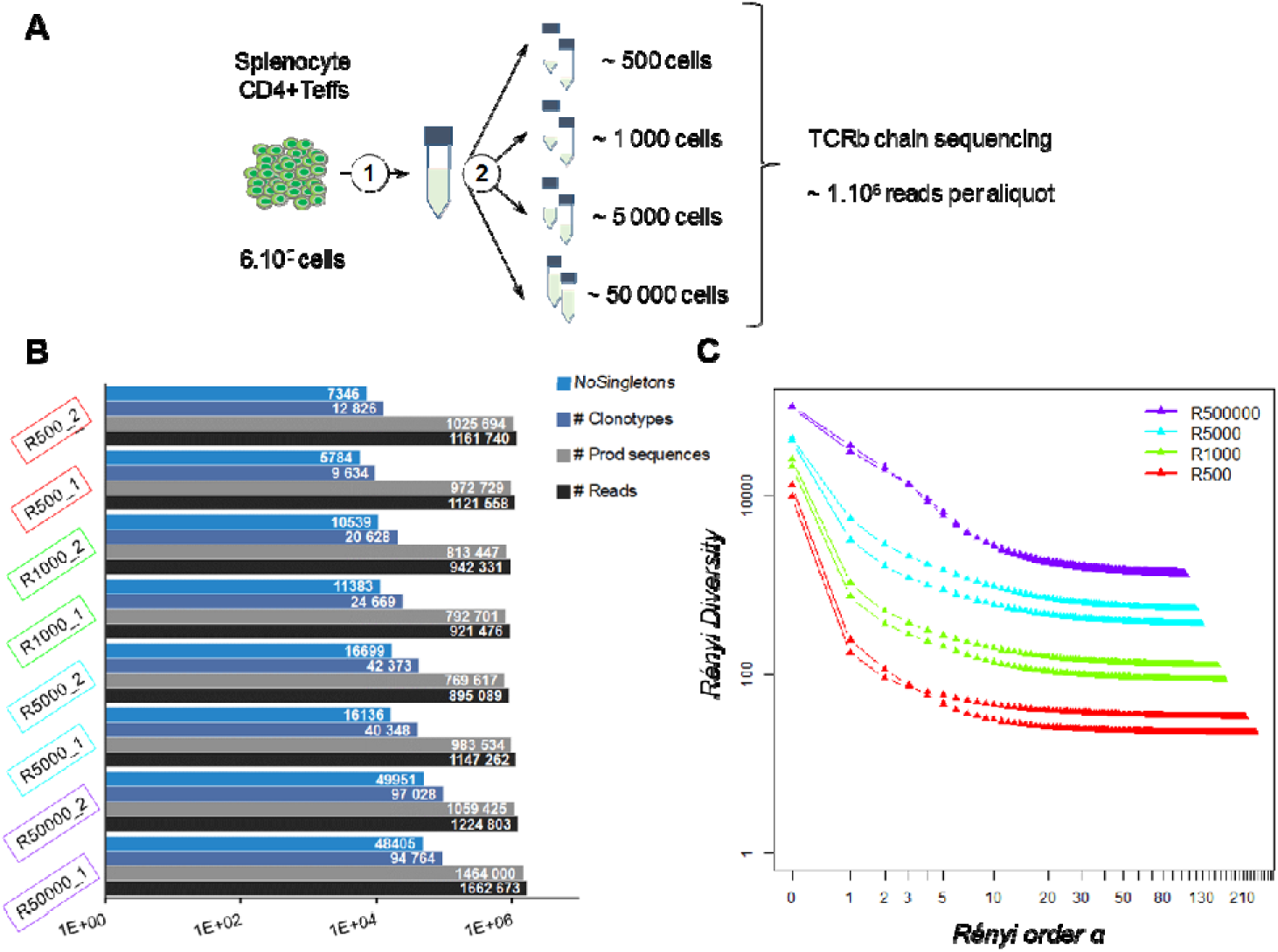
Impact of the sequencing depth on the measured diversity. **A: Experimental design:** 600 000 CD4+Foxp3-cells were sorted from female Balb/C Foxp3 splenocytes. RNA was extracted (1) and split into aliquots equivalent to the amount of mRNA of 500, 1 000, 5 000 and 50 000 cells (2). Two aliquots were produced for each amount of RNA. The eight prepared aliquots were processed for TRB chain deep sequencing. **B: Dataset summaries**. Histograms show, for each resulting dataset, the number of reads (black), productive TRB sequences (grey), observed clonotypes (blue) and clonotypes observed more than once (NoSingletons; light blue). **C: Diversity profiles**. For each dataset, a Rényi diversity profile was computed: diversity metrics using clonotype frequencies were calculated for increasing values of Rényi order α until stabilisation of the resulting diversity. For α = 1, the Shannon entropy was computed.

On average, 1.13 (+/− 0.16) million reads were produced for each aliquot (Supplemental Table I), which is in the average range of common practice (Greiff et al., 2015a; Mamedov et al., 2013; Rosati et al., 2017). As summarised in Figure 1B, 0.99.10^6^ (+/− 0.15.10^6^) TRB sequences were identified per aliquot regardless of the sample size. The point here is to determine whether the sample size will impact the resulting repertoire distribution.

**Table I.** Shannon diversity calculated for each dataset

| Shannon diversity | R500 | R1000 | R5000 | R50000 |
| --- | --- | --- | --- | --- |
| #1 | 171 | 1 034 | 3 124 | 30 432 |
| #2 | 238 | 735 | 5 337 | 35 027 |

Thus, we analysed the diversity of the observed repertoires according to sample size. It is noteworthy that the number of unique clonotypes (i.e. unique combination of TRBV - CDR3pep - TRBJ) per sample was always higher than the number of cells per sample. This discrepancy was more marked for small size samples, with approximately 20- to 2-fold more clonotypes per sample than cells with the “500-” and “50 000-cell” samples, respectively. In each dataset, about 50% (+/−6%) of the clonotypes were observed once (singletons). After removing the singletons, as is commonly done (Greiff et al., 2015a), this bias was reduced for the large samples, while the numbers of clonotypes remained much higher than the actual number of cells in small samples (Figure 1B). Still, overall richness remained equivalent between all sample sizes.

In order to refine the diversity assessment of these TRB repertoires, we computed their diversity profile (Figure 1C) applying Rényi entropy on the clonotype relative frequencies within each dataset. This function is used in ecological science to quantify the diversity, uncertainty and randomness of a given system (Ricotta, 2003; Schroeder, 2015). As the α order increases, it defines metrics spanning from (i) the species richness to (ii) the clonal dominance that progressively discard the scarcest species. The exponential of these metrics provides comparable effective numbers of species, used here to build a diversity profile. Analysis of the Rényi profiles for the eight aliquots showed that TRB repertoire diversity strongly decreases when the Rényi order α value increases. While richness was comparable between all sample sizes, diversity drops in proportion to sample size when progressively discarding scarce clonotypes to reach a plateau of clonotype counts below the initial number of cells.

### Shannon entropy as a threshold to filter the clonotypes

To avoid bias related to sample size, we normalised each dataset by randomly selecting 700 000 sequences, ranked the unique clonotypes from the most to the least predominant (clonotype rank) and plotted their abundance (clonotype count) to assess their distribution (Figure 2A). It is noteworthy that, while all the aliquots come from the same sample, the clonotype distributions within each dataset are different. The smaller a sample, the higher the most predominant clonotype counts, making it difficult to apply a filtering rule based on the count values. The Rényi profiles (Figure 1C) showed that the repertoire diversity collapses at a Rényi order α of 1, which corresponds to the Shannon diversity index (Rényi, 1961). Since the number of clonotypes assessed by the Shannon index (Table I) correlates best with sample size (Pearson coeff= 0.966, p-value = 9.62.10^−5^ and MH = 0.877 on original clonotype number and Pearson coeff= 0.995, p-value = 2.92.10^−7^ and MH = 0.996 after clonotype number determined by Shannon index), we chose to use this metric as a threshold to discard scarce “uninformative” clonotypes (SUC) that could result from experimental noise (shown in grey in Figure 2A) and keep only “informative” ones. As shown in Figure 2B, the clonotype relative distribution within each dataset is not significantly altered by this filtering. Interestingly, as shown in Figure 2C, this efficiently normalises values of the Piélou evenness index, a measure of clonotype evenness (Pielou, 1966), (filled squares) that otherwise strongly decreases when the clonotype number/cell number ratio increases in unfiltered datasets, revealing that too high a sequencing depth for small samples alters clonotype distributions (Figure 2C, empty circles).

**Figure 2:**
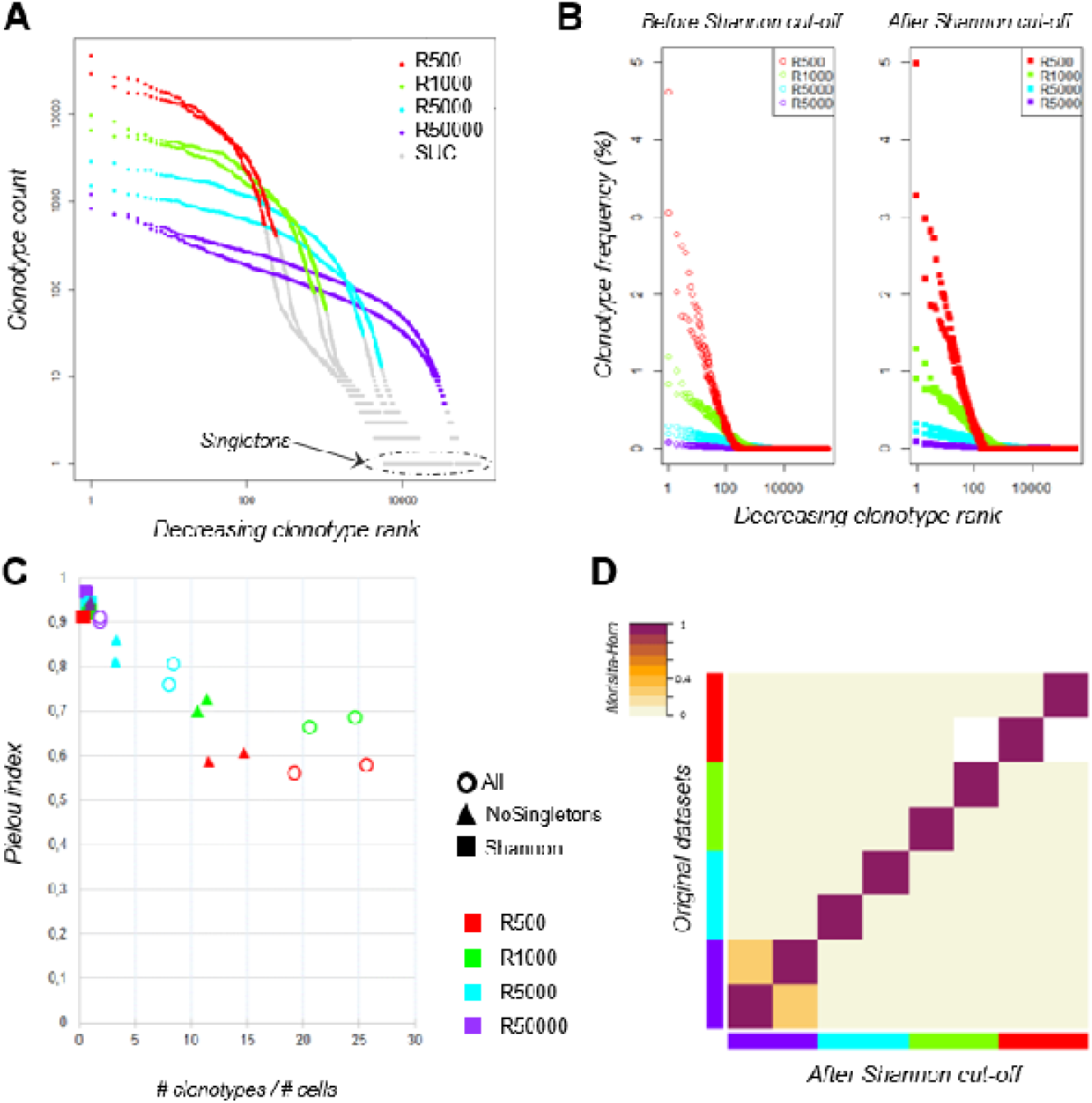
Clonotype distributions before and after data filtering. **A: TRB clonotype counts of the eight aliquots according to sampling size**. Within each dataset, clonotypes were ranked according to their counts from the most to the least predominant (decreasing clonotype rank) and their abundance (clonotype count). Both axes are log-scaled. Plots were coloured according to the sampling size: “500 cells” in red, “1 000 cells” in green, “5 000 cells” in cyan and “50 000 cells” in purple. Clonotypes filtered out using the Shannon index (see main text) are coloured in grey (SUC – Scarce Uninformative Clonotypes). **B: TRB clonotype distributions of the eight aliquots before and after data filtering**. Before (left) and after (right) filtering each dataset using the Shannon index as threshold, clonotypes were ranked from the most to the least predominant (decreasing clonotype rank) according to their relative frequencies (clonotype frequency). The X-axis is log-scaled. Distributions were coloured according to the sampling size as previously. **C: Impact of clonotype filtering on the clonotype distribution evenness**. The ratio between the number of clonotypes and the number of cells (x-axis) was calculated for each aliquot before (circles) and after clonotype filtering either by removing only singletons (triangles) or using the Shannon index as a threshold (squares). For each dataset, the Piélou evenness index was calculated (y-axis). Aliquots are identified according the sampling size as previously. **D: Similarity between datasets before and after Shannon filtering**. The Morisita-Horn similarity index between all pairs of datasets is colour-coded according to the indicated scale before (lower half-triangle) and after (upper half-triangle) Shannon filtering. Aliquots are identified according to sampling size as previously.

To confirm that the filtering does not bias the overall repertoire diversity, we computed the Morisita-Horn similarity index between the datasets before and after filtering; the high similarity values ([0.983; 0.997]) shown on the matrix diagonal in Figure 2D confirm that the datasets are not altered in the process. The similarity matrix also reveals a low similarity between replicates, except for the ‘50 000 cell’ samples, which are big enough to share rare clonotypes. Thus, high sequencing depth does not ensure good coverage of clonotype richness. This led us to question the robustness of RepSeq experiment results.

### Robustness of the TRB repertoire diversity assessment by RepSeq

We sorted 3.10^6^ Teff cells from splenocytes, extracted the RNA and split it into three equivalent RNA aliquots, and then sequenced them independently at a high depth targeting the TCRb chain using the iRepertoire^®^ multiplex PCR technology. On average, for each aliquot, 8.33 (+/− 0.66) million reads were produced and 5.63 (+/− 0.56) million TRB sequences were identified, among which an average of 130.10^3^ (+/− 5.10^3^) clonotypes (Supplemental Table II). After applying Shannon filtering, the dataset sizes were reduced to 4.7 (+/− 0.6) million TRB sequences for a total of 44 217 (+/− 304) clonotypes. Datasets were rarefied at an equivalent size by randomly selecting 4.10^6^ sequences for each sample.

We first analysed the clonotype distributions within each dataset. The three distributions were similar between replicates (Figure 3A). However, when we compared the composition of the three TRB repertoires by clonotype overlap, it appeared that about 36% of the clonotypes observed in each dataset are shared by another replicate, with only 6 599 clonotypes common to the three replicates. Although these shared clonotypes represent only 6% of the 105 332 clonotypes identified overall, their expression accounted for approximately 38% of each repertoire (Figure 3B). We then decomposed the clonotype collection by labelling the clonotypes as private (not shared between replicates) or shared by 2 or 3 replicates. For each dataset, clonotypes were sorted from the most to the least abundant and enrichment curves were built for each category according to the sharing status of each clonotype (Figure 3C). The resulting clonotype spectrum revealed that the most predominant clonotypes are shared by the three replicates, while the private clonotypes, which are the more numerous, are enriched for scarce clonotypes, therefore reducing the similarity between technical replicates. These results demonstrate that although the sampling of a large and polyclonal cell population has no impact on the observed clonotype distribution, the repertoire composition is affected: even if the most predominant clonotypes are always captured, a major proportion of the clonotypes observed with a single sequencing are private scarce ones. This observation confirms that the more abundant a clonotype, the more likely it is to be observed by sequencing. However, most rare clonotypes will remain unseen with a single sequencing run.

**Figure 3:**
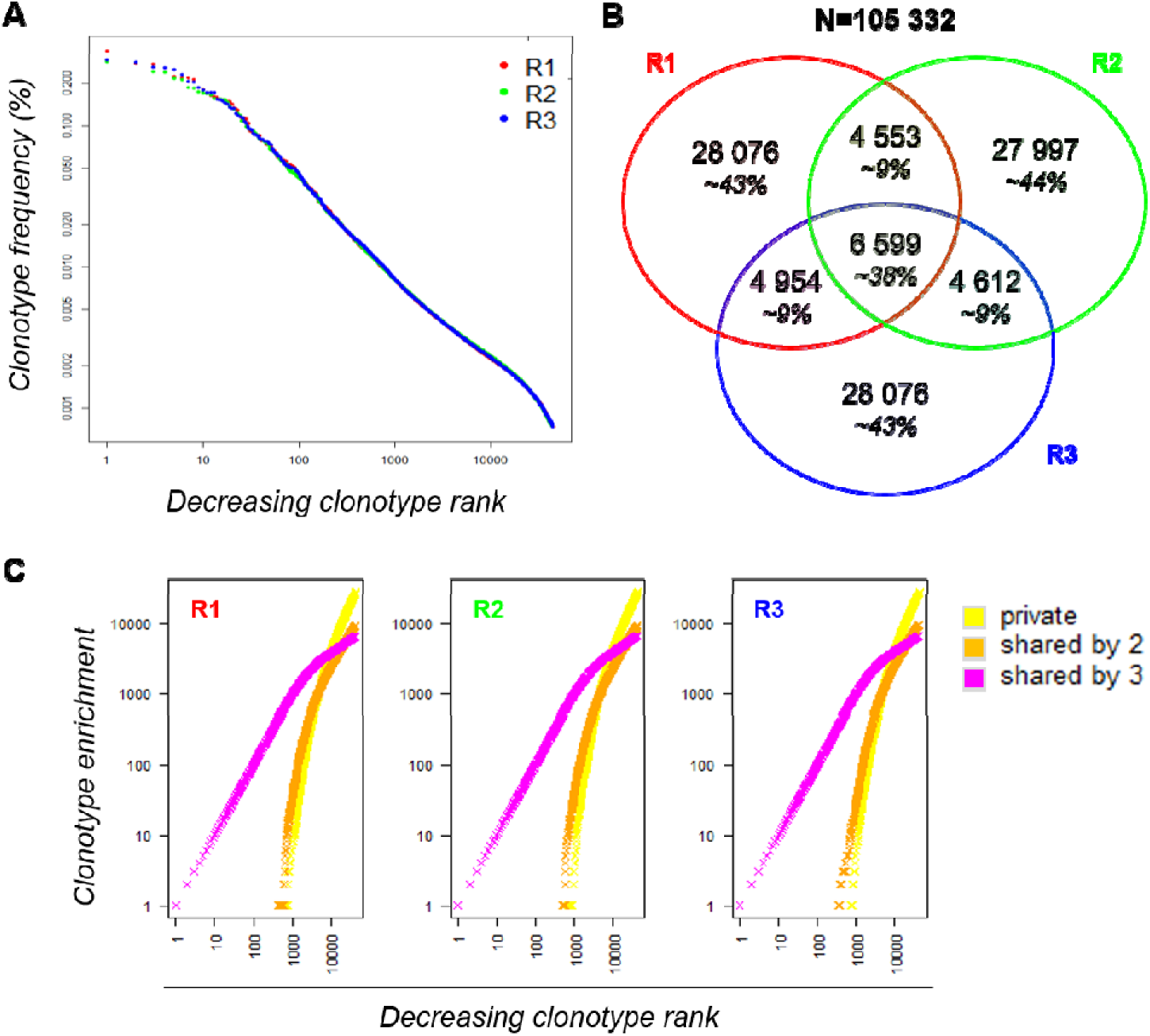
Robustness of a RepSeq experiment. **A: Clonotype distribution of the three replicates within each dataset**. Informative clonotypes were ranked decreasingly according to their abundance and their frequency plotted. The x axis is log-scaled. **B: Venn diagram between the three replicates**. Out of the 105 332 clonotypes observed in total, only 6 599 are shared by the three replicates; their cumulative frequency covers about 38% of each dataset. **C: Spectrum of unshared (yellow) and shared (by 2 in orange and by 3 in magenta) clonotypes in each replicate**. Within each dataset, clonotypes were ranked according to their counts from the most to the least predominant (decreasing clonotype rank). Since clonotypes are labelled according to their sharing status, the clonotype enrichment ((y-axis)) of each sharing group is incremented (+1) when a corresponding clonotype is found in the ranked list.

### Computational assessment of the impact of sequencing depth on observed diversity

In order to assess properly the representativeness of the diversity observed when analysing a clonotype repertoire by RepSeq, it would be necessary to know *a priori* its true diversity, which is not achievable with a classic experimental approach inherently subject to sampling bias.

Several studies have demonstrated that immune repertoires follow a Zipf-like distribution (Burgos and Moreno-Tovar, 1996; Greiff et al., 2015b; Mora and Walczak, 2016; Schwab et al., 2014; Sepúlveda et al., 2005), which translates a relation between rank order and frequency of occurrence: the frequency *f* of a particular observation is inversely proportional to its rank *r* (Aitchison et al., 2016) with:

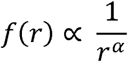

for Zipf-α parameter ≈ 1 (Piantadosi, 2014).

In addition, the lower the Zipf-α parameter of a distribution, the more evenly represented the clonotypes involved (Greiff et al., 2015b). We applied this observation to build clonotype distributions of a fixed size and known diversity to simulate the sampling effect occurring during a RepSeq experiment.

Seven Zipf distributions of 6.10^5^ sequences each were simulated with a parameter A=1/Zipf-α ranging from 2 to 100. These distributions were then assigned to a list of clonotypes randomly generated using the *TcR* package (Nazarov et al., 2015), leading to seven TRB clonotype repertoires of known diversity.

As observed in Figure 4A, the distribution slope varies according to the depth of sequencing of the clonotypes. For example, for the distribution simulated with A = 2 (A2), the resulting distribution is skewed in a way that clonotype counts range from 1 to 31 109, whereas when A = 100 (A100), clonotype counts do not exceed 9. These different distributions lead to datasets of varying richness, as summarised Table II.

**Table II.** Summary of the simulated Zipf distributions

| A | 2 | 3 | 4 | 5 | 10 | 20 | 100 |
| --- | --- | --- | --- | --- | --- | --- | --- |
| $\sum_{i=1}^{N_A} z_{A,i}$ | 6E+05 | 6E+05 | 6E+05 | 6E+05 | 6E+05 | 6E+05 | 6E+05 |
| $N_A$ | 155 495 | 394 784 | 435 528 | 450 625 | 469 974 | 476 829 | 480 919 |

**Figure 4:**
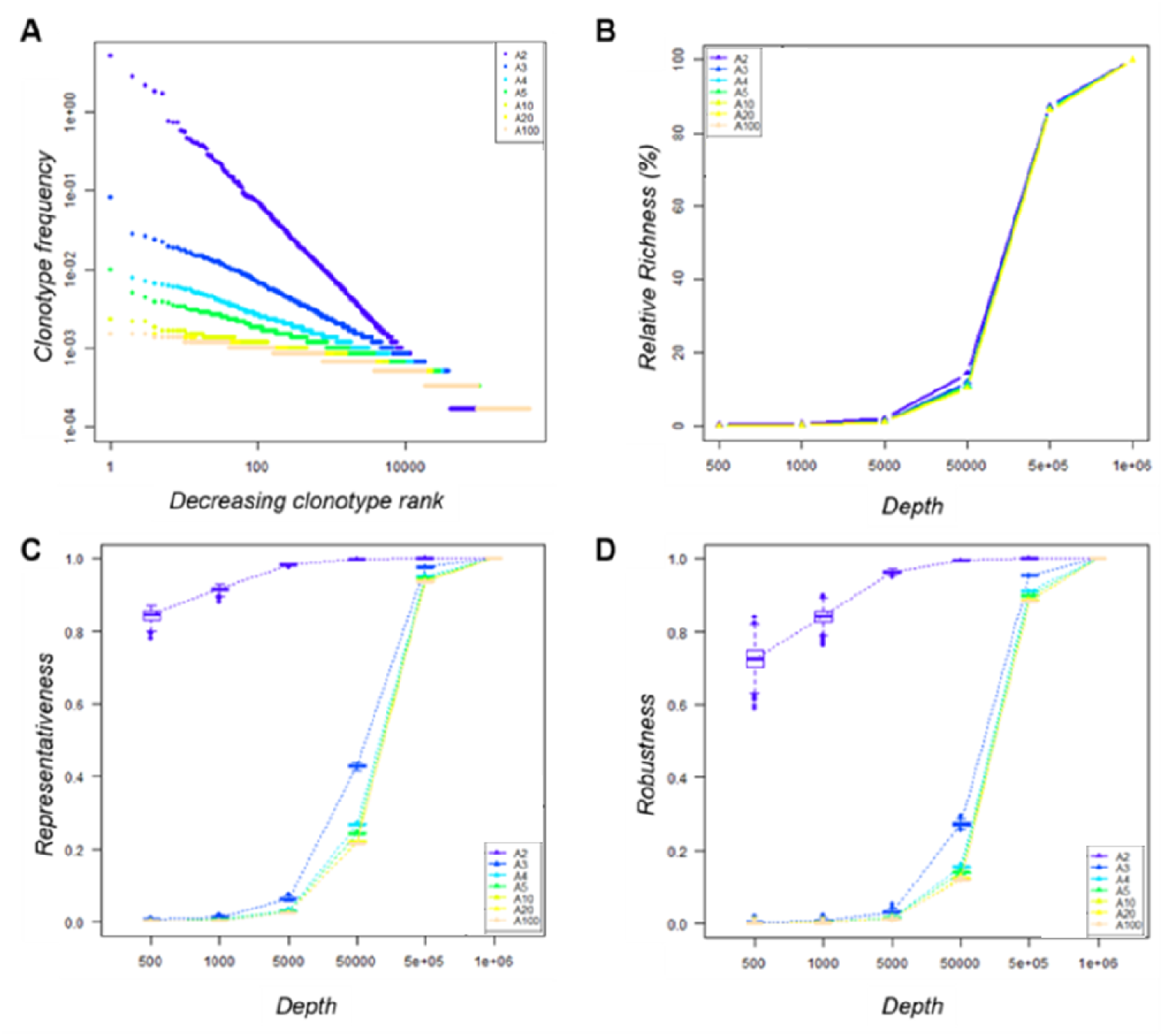
Impact of sequencing depth on the observed diversity. **A: Clonotype distribution within the seven simulated datasets** - Within each A-dataset, clonotypes were ranked decreasingly according to their abundance and their frequency was plotted. Both axes are log-scaled. Distributions are coloured according to the A parameter used to simulate it. **B: Impact of sequencing depth on the observed clonotype richness** - For a given A-dataset, clonotype richness was measured within the 100 subsamples produced for each depth and divided by that of the original dataset. The median value by depth is represented for each condition. The 95% CI was calculated but cannot be seen since it merged with the median value. **C: Representativeness of the sequencing** – The Morisita-Horn similarity index was calculated between each subsample and its original dataset. Boxplots across the 100 subsamples of a given depth are colour-coded according to the A condition. **D: Reproducibility of the sequencing** – For each A-dataset, the Morisita-Horn similarity index was calculated between paired subsamples of a given depth. Boxplots across the 100 subsamples of a given depth are colour-coded according to the A condition.

For each of our seven “known” repertoire distributions, we generated 100 subsamples at six sample size (from 500 to 1.10^6^ sequences) reflecting several levels of sequencing depth. The clonotype richness observed within each subsample increased according to the depth, as expected (Figure 4B). We used the Morisita-Horn similarity index to assess (i) representativeness (Figure 4C) by comparing the diversity captured for each subsample with the original repertoire diversity and (ii) reproducibility (Figure 4D) for the 100 subsamples for a given depth. When comparing the seven distributions at a given sequencing depth (5.10^4^ sequences, representing 8 % of the original repertoire), the representativeness of the diversity between distributions is different (Figure 4C), yet with similar relative richness values. For the ‘A2’ condition, the similarity index between this subsample and the original repertoire was above 0.8, while it varied from 0.2 to 0.5 for the other conditions (Figure 4C). A dataset of 5.10^5^ sequences (80% of the original repertoire size) is needed to reach a 0.9 similarity for the latter. However, a suitable representativeness does not ensure good reproducibility of the observations. With 500 or 1000 sequences, even if the diversity observed for the ‘A2’ condition is quite representative (MH ^~^ 0.8), the high variability between the subsamples implies a low reproducibility and thus an inability to observe exhaustively all the clonotypes (Figure 4D).

We sought to identify which simulated distribution would be the more representative of our experimental datasets. To this end, we compared the slope at the steepest descent point of each simulated distribution with those of all the experimental data analysed in this study. The experimental distribution slopes are most comparable with the ‘A3’ and ‘A5’ distributions, with the exception of that of the R500_2 sample (Supplemental Table III). Thus, we chose the ‘A3’ distribution dataset as the more representative. In order to understand the low overlap observed between experimental replicates in Figure 3B, for each size we compared the ‘A3’ simulated subsamples to determine the proportion of clonotypes shared by three independent subsamples, as performed experimentally in Figure 3. As summarised in Table III, the proportion of private and shared clonotypes varies according to the coverage of the initial repertoire stretch. For subsamples with sizes representing less than 1% of that of the initial dataset, almost all the clonotypes observed are private (only captured in one subsample). For the ‘5.10^4^ sequence’ subsamples, the size of which represents 8% of the original repertoire size, 16% of the clonotypes observed are captured at least twice. These proportions correspond to the observations we made in Figure 3 between the three experimental replicates. Finally, using subsamples of size close (80%) to that of the original, 95% of the observed clonotypes are shared by at least two replicates. In addition, as represented in Figure 5, at this depth, while one sample only captures about 12% of the overall existing clonotypes, three replicates cover a third of the overall richness. These observations suggest that multiple sequencing experiments can ensure greater clonotype exhaustiveness than a unique very deep sequencing.

**Figure 5:**
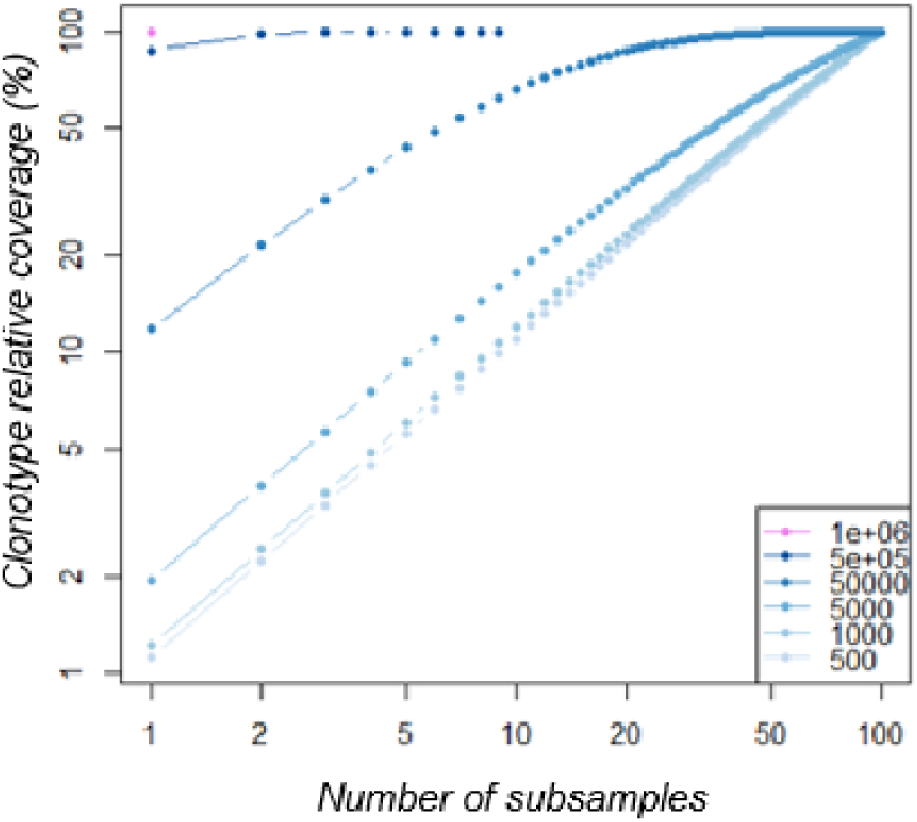
Clonotype coverage of A3-dataset richness increases with multiple subsamples. *The A3-datasets were subsampled at increasing depth (from 500 to 1.10*^6^ sequences as indicated in the legend from light to dark blue). For each depth, 100 subsamples were produced. Within each subsample series, an increasing number of subsamples (x-axis) were randomly selected and their cumulative clonotype richness was calculated relative to the original dataset richness (clonotype richness coverage).

**Table III.** Sharing proportion between three replicates

| Median proportion<br>of clonotypes observed | Dataset sizes |  |  |  |  |  |
| --- | --- | --- | --- | --- | --- | --- |
| | $5 \cdot 10^2$ | $1 \cdot 10^3$ | $5 \cdot 10^3$ | $5 \cdot 10^4$ | $5 \cdot 10^5$ | $1 \cdot 10^6$ |
| Private | 99.7 | 99.4 | 97.5 | 83.7 | 5.2 | - |
| Shared by 2 | 0.3 | 0.6 | 2.4 | 14.3 | 27.2 | - |
| Shared by 3 | - | - | 0.1 | 2 | 67.6 | 100 |
| Total number across 3 replicates | 1 493 | 2 973 | 14 456 | 117 634 | 393 434 | 394 784 |

## DISCUSSION

RepSeq offers new opportunities to identify biomarkers of health or disease by monitoring adaptive immune cell diversity at unprecedented high resolution. Continuing improvements in molecular biology protocols and sequencing technologies are increasing the accuracy of clonotype detection (Friedensohn et al., 2017). Still, clear evaluation of the reproducibility and representability of the observed diversity is missing. This is particularly true when considering small size samples, such as small cell subsets or cells from biopsies, though of utmost interest when studying TCR repertoires. Although over-sequencing has been recommended to ensure the identification of rare clonotypes (Mamedov et al., 2013), it does increase the risk of generating uninformative, possibly artefactual clonotypes such as duplicate reads and chimeric reads (Head et al., 2014). Indeed, when sequencing samples of varying sizes at a commonly used depth, we found that small datasets contained 20 times more clonotypes than what would be expected regarding to the sample size. This figure decreases when the starting material is increased, demonstrating that over-sequencing small samples dramatically generates noise that cannot be corrected by removing only singletons. Although the relationship between sample size and sequencing depth that we used may appear extreme, it can commonly occur when studying small cell subsets involved in immunological processes. Single-cell sequencing technologies can be an alternative, but may require more cells than actually recovered in particularly low input samples. These observations demonstrate the drawbacks of discarding clonotypes based only on their counts and the need for objective approaches in order to assess the actual richness of a repertoire effectively.

Here we provide a bioinformatics approach to assess accurately the number of unique clonotypes in a large and complex cell population, even when over-sequenced. When analysing the diversity profiles of repertoires from subsamples of varying sizes of a unique starting sample, we identified the Shannon entropy as a reliable threshold to eliminate clonotypes arising from technical noise (SUC) and to focus on informative TR clonotypes (Figure 1C, Figure 2A). This filtering strategy has no impact on the overall clonotype distribution (Figure 2B). Importantly, this approach was validated on subsamples originating from a single starting sample. Therefore, the representability of the smallest subsample was questioned. While the distribution evenness was sample size-dependent when considering all the reads, filtering by the Shannon entropy index removed this variability between replicates (Figure 2C). This proposed strategy therefore offers an accurate assessment of clonotype identification and representability, even in extreme situations.

Our results strongly suggest that sequencing depth must be adapted to the initial cell amount. We showed that “50 000 cell” replicates are closer to each other than lower input pairs of samples (Figure 2D). This observation emphasises the need to adapt the sample size to the population of interest. All aliquots analysed here were obtained from a rich and polyclonal cell population. In order to be reliable, a sample needs to be large enough to ensure that most of the clones are represented. Here, about 20% of the clonotypes observed in the two replicates (6 766 out of 30 422 and 35 020 clonotypes) are shared.

Altogether these results show how complex defining a RepSeq strategy can be in guaranteeing the representativeness of the repertoire diversity. If sequencing depth is not adapted to the population size, it can negatively affect the resulting observed diversity, in particular if data are not properly analysed. This is particularly crucial since the clonality of a population is rarely known before its sequencing, leading to misinterpretation of the results. Since the sequencing depth used was much higher than the size of the samples we analysed, one would expect good, if not exhaustive, coverage of the overall clonotypes. Conversely, we show that this is by no means the case, with only part of clonotypes being observed with confidence. These observations led us to question the robustness of the results of RepSeq experiments.

Multiple sequencing of the same sample revealed very low overlap between technical replicates, even after filtering out uninformative TR clonotypes, and merely captures the most frequent clonotypes. Rare clonotypes were at best shared by two replicates. As already suggested by Greiff et al. (Greiff et al., 2015a), our results are in favour of multiple sequencing when considering very diverse samples. This can be explained by the experimental sampling enforced by the different RepSeq steps (from RNA amplification to library sequencing). In order to validate these experimental observations and propose guidelines for RepSeq studies, we simulated different repertoire distributions and found that the representativeness of a very evenly distributed repertoire, which could be assimilated to a polyclonal repertoire, is more sensitive to the sequencing depth. The number of sequences produced (by multiple sequencing) needs to be equivalent to the population size to ensure a good assessment of the original diversity (Figure 4C). This is particularly true for small samples for which a too deep sequencing can favour the erroneous sequences possibly generated during library preparation (Heather et al., 2017) introducing experimental noise.

Altogether, we provide here a method to accurately discard uninformative clonotypes for small and large samples based on the application of Shannon diversity index threshold filtering, as well as guidelines for RepSeq experimental design. In addition, we show how computational simulation of diversity can improve adaptive repertoire analysis assessment where controlled reference repertoires with known actual diversity can be modelled and subject to experimental design and annotation tool flaws. We believe these will be useful in ensuring better RepSeq analyses when looking at rare or unknown cell populations participating in pathophysiological processes and will facilitate the discovery of HTS-based biomarkers.

## ACKNOWLEDGEMENTS

We are grateful to B. Gouritin for his help in cell sorting. We thank iRepertoire^®^ for providing us with the required data format to implement our analysis pipeline.

## REFERENCES

Aitchison, L., Corradi, N., and Latham, P.E. (2016). Zipf’s law arises naturally when there are underlying, unobserved variables. PLoS Comput. Biol. 12, e1005110.

Bashford-Rogers, R.J.M., Palser, A.L., Idris, S.F., Carter, L., Epstein, M., Callard, R.E., Douek, D. C., Vassiliou, G.S., Follows, G.A., Hubank, M., et al. (2014). Capturing needles in haystacks: a comparison of B-cell receptor sequencing methods. BMC Immunol. 15, 29.

Benichou, J., Ben-Hamo, R., Louzoun, Y., and Efroni, S. (2012). Rep-Seq: uncovering the immunological repertoire through next-generation sequencing: Rep-Seq: NGS for the immunological repertoire. Immunology 135,183–191.

Bergot, A.-S., Chaara, W., Ruggiero, E., Mariotti-Ferrandiz, E., Dulauroy, S., Schmidt, M., von Kalle, C., Six, A., and Klatzmann, D. (2015). TCR sequences and tissue distribution discriminate the subsets of naïve and activated/memory Treg cells in mice: Molecular immunology. Eur. J. Immunol. 45, 1524–1534.

Boudinot, P., Mariotti-Ferrandiz, M.E., Pasquier, L.D., Benmansour, A., Cazenave, P.-A., and Six, A. (2008). New perspectives for large-scale repertoire analysis of immune receptors. Mol. Immunol. 45, 2437–2445.

Burgos, J.D., and Moreno-Tovar, P. (1996). Zipf-scaling behavior in the immune system. Biosystems 39, 227–232.

Dash, P., Fiore-Gartland, A.J., Hertz, T., Wang, G.C., Sharma, S., Souquette, A., Crawford, J.C., Clemens, E.B., Nguyen, T.H.O., Kedzierska, K., et al. (2017). Quantifiable predictive features define epitope-specific T cell receptor repertoires. Nature 547, 89–93.

Davis, M.M., and Bjorkman, P.J. (1988). T-cell antigen receptor genes and T-cell recognition. Nature 334, 395–402.

Dong, M., Artusa, P., Kelly, S.A., Fournier, M., Baldwin, T.A., Mandl, J.N., and Melichar, H.J. (2017). Alterations in the Thymic Selection Threshold Skew the Self-Reactivity of the TCR Repertoire in Neonates. J. Immunol. 199, 965–973.

Emerson, R.O., DeWitt, W.S., Vignali, M., Gravley, J., Hu, J.K., Osborne, E.J., Desmarais, C., Klinger, M., Carlson, C.S., Hansen, J.A., et al. (2017). Immunosequencing identifies signatures of cytomegalovirus exposure history and HLA-mediated effects on the T cell repertoire. Nat. Genet. 49, 659–665.

Evert, S., and Baroni, M. (2007). zipfR: Word frequency distributions in R. In Proceedings of the 45th Annual Meeting of the ACL on Interactive Poster and Demonstration Sessions, (Association for Computational Linguistics), pp. 29–32.

Fisher, R.A., Steven-Corbet, A., and Williams, C.B. (1943). The relation between the number of species and the number of individuals in a random sample of an animal population. J. Anim. Ecol. 12, 42–58.

Freeman, J.D., Warren, R.L., Webb, J.R., Nelson, B.H., and Holt, R.A. (2009). Profiling the T-cell receptor beta-chain repertoire by massively parallel sequencing. Genome Res. 19, 1817–1824.

Friedensohn, S., Khan, T.A., and Reddy, S.T. (2017). Advanced Methodologies in High-Throughput Sequencing of Immune Repertoires. Trends Biotechnol. 35, 203–214.

Glanville, J., Huang, H., Nau, A., Hatton, O., Wagar, L.E., Rubelt, F., Ji, X., Han, A., Krams, S.M., Pettus, C., et al. (2017). Identifying specificity groups in the T cell receptor repertoire. Nature.

Greiff, V., Miho, E., Menzel, U., and Reddy, S.T. (2015a). Bioinformatic and Statistical Analysis of Adaptive Immune Repertoires. Trends Immunol. 36, 738–749.

Greiff, V., Bhat, P., Cook, S.C., Menzel, U., Kang, W., and Reddy, S.T. (2015b). A bioinformatic framework for immune repertoire diversity profiling enables detection of immunological status. Genome Med. 7.

Head, S.R., Komori, H.K., LaMere, S.A., Whisenant, T., Van Nieuwerburgh, F., Salomon, D.R., and Ordoukhanian, P. (2014). Library construction for next-generation sequencing: Overviews and challenges. BioTechniques 56.

Heather, J.M., Best, K., Oakes, T., Gray, E.R., Roe, J.K., Thomas, N., Friedman, N., Noursadeghi, M., and Chain, B. (2016). Dynamic Perturbations of the T-Cell Receptor Repertoire in Chronic HIV Infection and following Antiretroviral Therapy. Front. Immunol. 6.

Heather, J.M., Ismail, M., Oakes, T., and Chain, B. (2017). High-throughput sequencing of the T-cell receptor repertoire: pitfalls and opportunities. Brief. Bioinform.

van Heijst, J.W.J., Ceberio, I., Lipuma, L.B., Samilo, D.W., Wasilewski, G.D., Gonzales, A.M.R., Nieves, J.L., van den Brink, M.R.M., Perales, M.A., and Pamer, E.G. (2013). Quantitative assessment of T cell repertoire recovery after hematopoietic stem cell transplantation. Nat. Med. 19, 372–377.

Hill, M.O. (1973). Diversity and evenness: a unifying notation and its consequences. Ecology 54, 427–432.

Horn, H.S. (1966). Measurement of “Overlap” in Comparative Ecological Studies. Am. Nat. 100, 419–424.

Kuang, M., Cheng, J., Zhang, C., Feng, L., Xu, X., Zhang, Y., Zu, M., Cui, J., Yu, H., Zhang, K., et al. (2017). A novel signature for stratifying the molecular heterogeneity of the tissue-infiltrating T-cell receptor repertoire reflects gastric cancer prognosis. Sci. Rep. 7, 7762.

Lai, L., Wang, L., Chen, H., Zhang, J., Yan, Q., Ou, M., Lin, H., Hou, X., Chen, S., Dai, Y., et al. (2016). T cell repertoire following kidney transplantation revealed by high-throughput sequencing. Transpl. Immunol. 39, 34–45.

Langerak, A.W., Brüggemann, M., Davi, F., Darzentas, N., van Dongen, J.J.M., Gonzalez, D., Cazzaniga, G., Giudicelli, V., Lefranc, M.-P., Giraud, M., et al. (2017). High-Throughput Immunogenetics for Clinical and Research Applications in Immunohematology: Potential and Challenges. J. Immunol. 198, 3765–3774.

Levenshtein, V.I. (1966). Binary codes capable of correcting deletions, insertions, and reversals. Sov. Phys. Dokl. 10, 707–710.

Maceiras, A.R., Almeida, S.C.P., Mariotti-Ferrandiz, E., Chaara, W., Jebbawi, F., Six, A., Hori, S., Klatzmann, D., Faro, J., and Graca, L. (2017). T follicular helper and T follicular regulatory cells have different TCR specificity. Nat. Commun. 8, 15067.

Madi, A., Shifrut, E., Reich-Zeliger, S., Gal, H., Best, K., Ndifon, W., Chain, B., Cohen, I.R., and Friedman, N. (2014). T-cell receptor repertoires share a restricted set of public and abundant CDR3 sequences that are associated with self-related immunity. Genome Res. 24, 1603–1612.

Madi, A., Poran, A., Shifrut, E., Reich-Zeliger, S., Greenstein, E., Zaretsky, I., Arnon, T., Van Laethem, F., Singer, A., Lu, J., et al. (2017). T cell receptor repertoires of mice and humans are clustered in similarity networks around conserved public CDR3 sequences. ELife 6.

Magurran, A. (2004). Measuring Biological Diversity (Wiley).

Mamedov, I.Z., Britanova, O.V., Zvyagin, I.V., Turchaninova, M.A., Bolotin, D.A., Putintseva, E. V., Lebedev, Y.B., and Chudakov, D.M. (2013). Preparing Unbiased T-Cell Receptor and Antibody cDNA Libraries for the Deep Next Generation Sequencing Profiling. Front. Immunol. 4.

Mariotti-Ferrandiz, E., Pham, H.-P., Dulauroy, S., Gorgette, O., Klatzmann, D., Cazenave, P.-A., Pied, S., and Six, A. (2016). A TCRβ Repertoire Signature Can Predict Experimental Cerebral Malaria. PLOS ONE 11, e0147871.

Marrero, I., Hamm, D.E., and Davies, J.D. (2013). High-Throughput Sequencing of Islet-Infiltrating Memory CD4+ T Cells Reveals a Similar Pattern of TCR VÎ^2^ Usage in Prediabetic and Diabetic NOD Mice. PLoS ONE 8, e76546.

Marrero, I., Aguilera, C., Hamm, D.E., Quinn, A., and Kumar, V. (2016). High-throughput sequencing reveals restricted TCR Vβ usage and public TCRβ clonotypes among pancreatic lymph node memory CD4+ T cells and their involvement in autoimmune diabetes. Mol. Immunol. 74, 82–95.

Mora, T., and Walczak, A. (2016). Quantifying lymphocyte receptor diversity. ArXiv Prepr. ArXiv160400487.

Murugan, A., Mora, T., Walczak, A.M., and Callan, C.G. (2012). Statistical inference of the generation probability of T-cell receptors from sequence repertoires. Proc. Natl. Acad. Sci. 109, 16161–16166.

Nazarov, V.I., Pogorelyy, M.V., Komech, E.A., Zvyagin, I.V., Bolotin, D.A., Shugay, M., Chudakov, D.M., Lebedev, Y.B., and Mamedov, I.Z. (2015). tcR: an R package for T cell receptor repertoire advanced data analysis. BMC Bioinformatics 16.

Oksanen, J., Blanchet, F.G., Kindt, R., Legendre, P., Minchin, P.R., O’hara, R.B., Simpson, G.L., Solymos, P., Stevens, M.H.H., and Wagner, H. (2013). Package ‘vegan.’ Community Ecol. Package Version 2.

Piantadosi, S.T. (2014). Zipf’s word frequency law in natural language: A critical review and future directions. Psychon. Bull. Rev. 21, 1112–1130.

Pielou, E.C. (1966). The measurement of diversity in different types of biological collections. J.Theor. Biol. 13, 131–144.

Plessy, C., Mariotti-Ferrandiz, E., Manabe, R.-I., and Hori, S. (2015). clonotypeR - high throughput analysis of T cell antigen receptor sequences. Biorxiv.

Poschke, I., Flossdorf, M., and Offringa, R. (2016). Next-generation TCR sequencing - a tool to understand T-cell infiltration in human cancers. J. Pathol. 240, 384–386.

Pugliese, A. (2017). Autoreactive T cells in type 1 diabetes. J. Clin. Invest. 127, 2881–2891.

Rényi, A. (1961). On measures of entropy and information. Proc. Fourth Berkeley Symp. Math. Stat. Probab. Univ. Calif. Press 1, 547–561.

Ricotta, C. (2003). On parametric evenness measures. J. Theor. Biol. 222, 189–197.

Robins, H.S., Srivastava, S.K., Campregher, P.V., Turtle, C.J., Andriesen, J., Riddell, S.R., Carlson, C.S., and Warren, E.H. (2010). Overlap and effective size of the human CD8+ T-cell receptor repertoire. Sci. Transl. Med. 2, 47ra64.

Rosati, E., Dowds, C.M., Liaskou, E., Henriksen, E.K.K., Karlsen, T.H., and Franke, A. (2017). Overview of methodologies for T-cell receptor repertoire analysis. BMC Biotechnol. 17.

Rossetti, M., Spreafico, R., Consolaro, A., Leong, J.Y., Chua, C., Massa, M., Saidin, S., Magni-Manzoni, S., Arkachaisri, T., Wallace, C.A., et al. (2017). TCR repertoire sequencing identifies synovial Treg cell clonotypes in the bloodstream during active inflammation in human arthritis. Ann. Rheum. Dis. 76, 435–441.

Schroeder, H.W. (2015). The Evolution and Development of the Antibody Repertoire. Front. Immunol. 6.

Schwab, D.J., Nemenman, I., and Mehta, P. (2014). Zipf’s Law and Criticality in Multivariate Data without Fine-Tuning. Phys. Rev. Lett. 113.

Seay, H.R., Yusko, E., Rothweiler, S.J., Zhang, L., Posgai, A.L., Campbell-Thompson, M., Vignali, M., Emerson, R.O., Kaddis, J.S., Ko, D., et al. (2016). Tissue distribution and clonal diversity of the T and B cell repertoire in type 1 diabetes. JCI Insight 1.

Sepúlveda, N., Boucontet, L., Pereira, P., and Carneiro, J. (2005). Stochastic Modeling of T cell receptor gene rearrangement. J. Theor. Biol. 234, 153–165.

Shugay, M., Bolotin, D.A., Putintseva, E.V., Pogorelyy, M.V., Mamedov, I.Z., and Chudakov, D.M. (2013). Huge Overlap of Individual TCR Beta Repertoires. Front. Immunol. 4.

Sims, J.S., Grinshpun, B., Feng, Y., Ung, T.H., Neira, J.A., Samanamud, J.L., Canoll, P., Shen, Y., Sims, P.A., and Bruce, J.N. (2016). Diversity and divergence of the glioma-infiltrating T-cell receptor repertoire. Proc. Natl. Acad. Sci. 113, E3529–E3537.

Six, A., Mariotti-Ferrandiz, M.E., Chaara, W., Magadan, S., Pham, H.-P., Lefranc, M.-P., Mora, T., Thomas-Vaslin, V., Walczak, A.M., and Boudinot, P. (2013). The past, present, and future of immune repertoire biology - the rise of next-generation repertoire analysis. Front. Immunol. 4, 413.

Thapa, D.R., Tonikian, R., Sun, C., Liu, M., Dearth, A., Petri, M., Pepin, F., Emerson, R.O., and Ranger, A. (2015). Longitudinal analysis of peripheral blood T cell receptor diversity in patients with systemic lupus erythematosus by next-generation sequencing. Arthritis Res. Ther. 17.

Theil, A., Wilhelm, C., Kuhn, M., Petzold, A., Tuve, S., Oelschlägel, U., Dahl, A., Bornhäuser, M., Bonifacio, E., and Eugster, A. (2017). T cell receptor repertoires after adoptive transfer of expanded allogeneic regulatory T cells: T cell receptor repertoires post-T_reg_ cell therapy. Clin. Exp. Immunol. 187, 316–324.

Thomas, N., Best, K., Cinelli, M., Reich-Zeliger, S., Gal, H., Shifrut, E., Madi, A., Friedman, N., Shawe-Taylor, J., and Chain, B. (2014). Tracking global changes induced in the CD4 T-cell receptor repertoire by immunization with a complex antigen using short stretches of CDR3 protein sequence. Bioinformatics 30, 3181–3188.

Warren, R.L., Nelson, B.H., and Holt, R.A. (2009). Profiling model T-cell metagenomes with short reads. Bioinformatics 25, 458–464.

Zhao, Y., Nguyen, P., Ma, J., Wu, T., Jones, L.L., Pei, D., Cheng, C., and Geiger, T.L. (2016). Preferential Use of Public TCR during Autoimmune Encephalomyelitis. J. Immunol. 196, 4905–4914.

